# A graphical approach of the interplay of eco-evolutionary dynamics and coexistence

**DOI:** 10.64898/2026.02.06.704293

**Authors:** Nicolas Loeuille, Rudolf P. Rohr

**Affiliations:** Sorbonne Université, UPEC, CNRS, IRD, INRA, Institute of Ecology and Environmental Sciences, IEES, F-75005 Paris, France; Department of Biology — Ecology and Evolution, University of Fribourg, Chemin du Musée 15, CH-1700 Fribourg, Switzerland

**Keywords:** SPECIES RESISTANCE, STRUCTURAL STABILITY, ADAPTIVE DYNAMICS

## Abstract

Given the accumulation of evidence that evolution can affect ecological dynamics, especially under global change scenarios, a key question is how such ecoevolutionary dynamics may change the coexistence of species and biodiversity in general. In the present article, we propose a graphical approach allowing to simultaneously discuss ecological coexistence and phenotype evolution. Our graphical approach allows tackling the two aspects in the same parameter space, allowing direct links between ecological and evolutionary perspectives. While evolution is often thought positive for the resilience of ecological systems, we first highlight it does not usually allow for better coexistence for the system as a whole. Even when focusing on the fate of the species that is evolving, evolution often leads to greater vulnerability. The graphical approach we propose is flexible and can be applied to all interaction types and covers variations in trade-off structures. Using this flexibility, we highlight how evolutionary effects can be positive or negative for coexistence, depending on these two components. Finally, we illustrate how the approach can be applied, using empirical examples derived from the literature. We thereby highlight the critical ingredients needed to inform the graphical approach, its potential use for proposing testable scenarios, but also clarify its limits.

## 1 Introduction

Traditionally, the maintenance of species diversity within ecological communities has been explained through a combination of abiotic constraints (environmental filtering, Kraft et al. (2015)) and biotic interactions among local species (e.g. competitive exclusion, Thorn et al. (2016), intransitive competition Vandermeer and Perfecto (2023), vertical food-chain assembly Hairston et al. (1960); Oksanen et al. (1981)), two processes that are often difficult to disentangle (Cadotte and Tucker, 2017). While these ecological components provide a fundamental basis for coexistence, recent works have stressed how evolution can also directly affect ecological dynamics (Hairston Jr et al., 2005; Carroll et al., 2014). Such evolutionary dynamics alter local interactions and environmental responses, thereby reshaping coexistence conditions. Understanding how eco-evolutionary dynamics influence community assembly has therefore emerged as a key question (Edwards et al., 2018; Leibold et al., 2022), particularly in the context of global changes. With communities increasingly exposed to disturbances and reorganization (Tylianakis et al., 2008), and with multiple documented cases of evolution in this context (Bonnet et al., 2022), assessing the consequences of evolution for biodiversity remains a central challenge (Urban et al., 2016).

At first sight, evolution may be expected to favour diversity. On macroevolutionary scales, it has produced Earth’s impressive biological richness through repeated speciation events. On shorter timescales, following disturbances, natural selection may promote adapted phenotypes, reversing population declines and preventing local extinctions — a process known as evolutionary rescue (Gomulkiewicz and Holt, 1995). By fostering adaptation, evolution could therefore buffer diversity loss. However, this perspective relies on the assumption that evolution generally increases the average growth rate of populations, an assumption that may not hold across ecological contexts (Dieckmann and Ferrière, 2004; Metz et al., 2008; Rohr and Loeuille, 2023). In fact, evolution can reduce abundances and even drive extinctions, both for the evolving population (evolutionary deterioration (Matsuda and Abrams, 1994), evolutionary suicide (Parvinen, 2005)), and at the community scale, where adapted species may exert stronger ecological pressures on others (Loeuille, 2019) or when evolution directly weakens ecological interactions (Weinbach et al., 2022). The overall role of evolution in maintaining diversity thus remains ambiguous and requires closer examination.

The consequences of evolution for coexistence depend critically on two aspects. First, the type of interaction under selection strongly shapes outcomes. In competitive contexts, evolution may promote niche displacement (Doebeli and Dieckmann, 2000; Lepori et al., 2024) or emergent neutrality (Scheffer and Van Nes, 2006), thereby favouring coexistence. In mutualistic interactions, evolution may either strengthen the interaction—stabilizing coexistence — or weaken it, undermining persistence (Toby Kiers et al., 2010; Weinbach et al., 2022; Yacine and Loeuille, 2024). In antagonistic interactions, adaptive foraging by predators can generate negative frequency dependence, favouring rare prey and thus maintaining diversity (Kondoh, 2003; Loeuille, 2010*a*). Trophic structure is also sensitive to evolutionary change: for example, the evolution of plant defences can limit energy transfer along food chains, constraining the number of trophic levels (Loeuille and Loreau, 2004; Koffel et al., 2018; Leibold et al., 2022). Next to ecological aspects, evolutionary constraints such as trade-offs are equally important. Traits that incur allocative costs (e.g. diverting resources from maintenance or reproduction, Herms and Mattson (1992)) may reduce population densities and energy availability for higher trophic levels. Traits can also generate ecological costs (Strauss et al., 2002; Müller-Schärer et al., 2004), where improving one interaction (e.g. stronger mutualism) simultaneously exacerbates another (e.g. greater susceptibility to herbivory). Evolution of such traits then creates immediate indirect ecological effects, with substantial consequences for coexistence.

Recent advances in coexistence theory, especially in the structural approach, offer new tools to study species maintenance through indices assessing the potential of coexistence and the resistance of each species to environmental perturbations, within a tractable but generalizable framework (Rohr et al., 2014; Saavedra et al., 2017; Medeiros et al., 2021; Lepori et al., 2024). This approach partitions coexistence across scales: at the community level, the *solid angle* quantifies the overall potential for coexistence, while at the species level, a *resistance angle* measure indicates how close a species is to its persistence threshold. Both depend on growth rates and interaction coefficients, which define the range of conditions under which communities persist. Because evolution directly alters growth and interaction rates, it shifts coexistence boundaries in predictable ways. In simple systems, these concepts can be illustrated graphically, making the framework accessible and intuitive. These illustrations are based on a geometric space defined by species growth rates, in which vectors embodying interactions are displayed ((Rohr et al., 2014; Saavedra et al., 2017). Because growth rates and interaction vectors are also directly linked to benefits and costs of phenotypic traits, the graphical approach can also be used to discuss the likely direction of evolution, thereby linking eco-evolutionary dynamics and coexistence in a unified geometrical space.

The aim of this article is to provide such graphical tools for exploring the interplay between evolution and coexistence. Our framework integrates ecological (structural stability) and evolutionary (adaptive dynamics) perspectives within a unified approach. We demonstrate its flexibility by applying it to diverse ecological interactions and tradeoffs. Our results reveal that evolutionary effects on coexistence are not systematically positive, even for the evolving species, but instead depend on the type of interaction and trade-off at hand. Thus, no universal answer exists — an unsurprising outcome given the complexity of eco-evolutionary processes. Nevertheless, the graphical approach (i) clarifies the conditions under which evolution is likely to have positive or negative effects, and (ii) highlights the empirical information most critical for evaluating these outcomes. After outlining the method, we illustrate its applicability with two case studies: (i) the evolution of plant defences and its consequences on plant–herbivore communities (based on Müller-Schärer et al. (2004)), and (ii) the well-documented role of body size in shaping growth rates and interaction strengths in competitive and predatory contexts.

## 2 Geometric approach

We build our geometric approach of the link between evolution and coexistence in two steps. First, following earlier articles on structural stability (Rohr et al., 2014; Saavedra et al., 2017), we show potential coexistence based on the growth rates of the two species, given their interactions (figure 1A). Second, we highlight that vector length and position may help to predict the most likely direction of evolution (e.g. selection or counter-selection of traits linked to *α*_11_ on figure 1B and 1C respectively), based on adaptive dynamics techniques. Once this likely direction is found out, implications of evolution for various aspects of coexistence can be analysed (arrows on Figure 1B and 1C). Importantly, our analysis reveals that the direction of natural selection and the effects of adaptive dynamics are intimately linked to the structural stability geometry, which allows us to assemble the two frameworks in a single geometrical approach.

**Figure 1:**
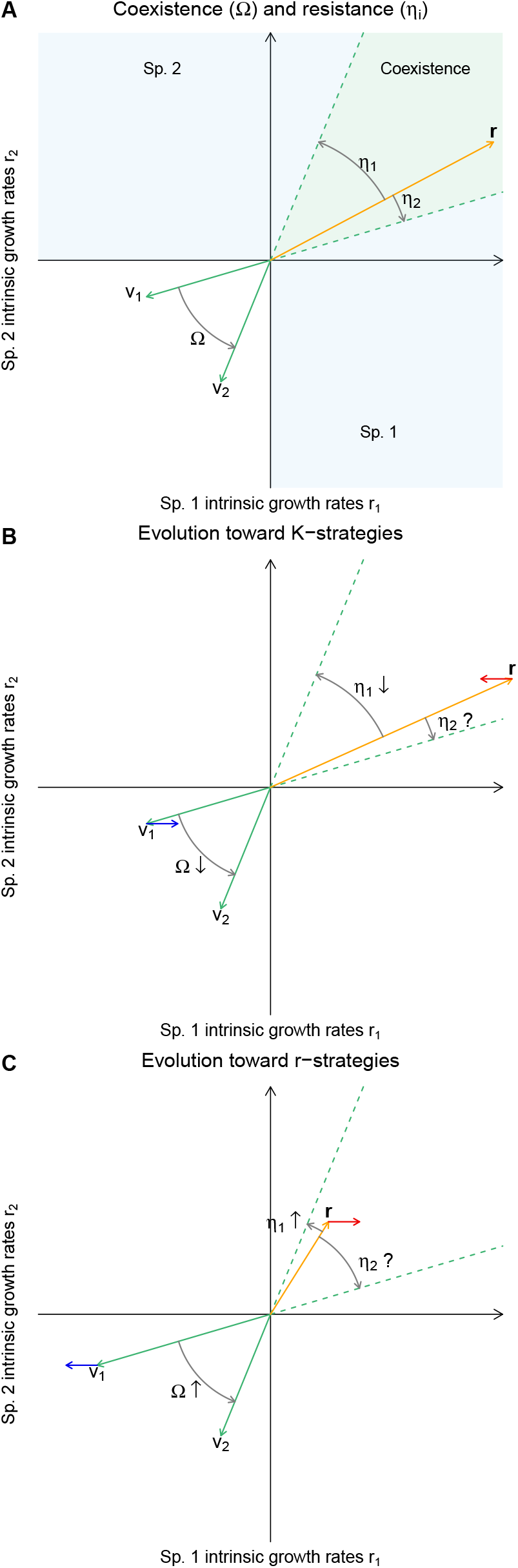
Combining the geometric approach to coexistence and to evolution. Panel **A** illustrates the community-level overall coexistence potential Ω and the species-level resistance *η*_*i*_ metrics of coexistence for a system of two competing species. In the 2-dimensional space made by species 1 and 2 intrinsic growth rates (*r*_1_ and *r*_2_), coexistence require that the vector of intrinsic growth rates **r** is inside the green cone generated by vectors *v*_1_ and *v*_2_ defined by ecological interactions. Therefore, the angles *η*_1_ and *η*_2_ between **r** and the border of green cone quantify species resistance, i.e. the perturbation needed to lose the species, while the angle Ω quantifies the overall potential of coexistence. (**B-C**) Vectors and angles also provide the likely direction of evolution (derivation in the text). On panel **B**, species 1 is likely to evolve toward K-strategies (*α*_11_ increases (blue arrow) at the expense of intrinsic growth rate (red arrow). Effects of evolution on coexistence metrics are then reported using arrows next to coexistence components Ω, *η*_1_ and *η*_2_. Panel **C** conversely shows a case where evolution favours r-strategies.

### 2.1 Coexistence

More precisely, let us start by presenting how panel A of figure 1 is constructed. Consider a (general) Lotka-Volterra model of two interacting species (Volterra, 1931; Case, 2000):

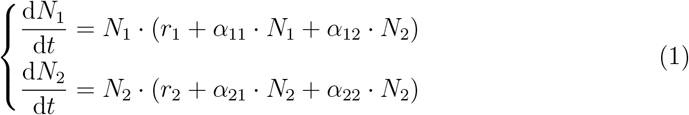

The intrinsic growth rate of species *i* is noted *r*_*i*_, while *α*_*ii*_ corresponds to its intraspecific competition, assumed to be negative. Parameters *α*_*ij*_ correspond to the ecological effects of species *j* on species *i*, i.e. per capita interaction effects, and their signs determine the types of interactions (antagonistic, competitive, or mutualistic).

Figure 1A shows how the conditions leading to the coexistence of two species in competition can be represented geometrically (MacArthur, 1972; Vandermeer, 1975; Pielou, 1977; Logofet, 1993; Saavedra et al., 2017). Briefly, the interaction coefficients *α*_*ij*_ determine the two vectors **v**_1_ = [*α*_11_, *α*_2_1]^*t*^ (quantifying ecological effects of species 1 on itself and on the other species) and **v**_2 =_ [*α*_11_, *α*_21_]^*t*^ (ecological effects of species 2). In turn, these vectors determine the so-called domain of feasibility (green cone on Fig1A), i.e. the set of intrinsic growth rates **r =**[*r*_1_, *r*_2_]^*t*^, leading to the existence of a positive equilibrium. Whenever vector **r** is inside the green cone, there exists an equilibrium at which both species have strictly positive abundances. To get coexistence, one needs stability on top of the feasibility, which is ensured whenever *α*_11_ · *α*_22_ − *α*_12_ · *α*_21_ *>* 0 (MacArthur and Levins, 1967; Case, 2000). This inequality can also be geometrically represented by the directed angle between the vectors **v**_1_ and **v**_2_, and this angle called Ω, must then fulfil 0^°^ *<* Ω *<* 180^°^. The angle Ω also defines the opportunity for feasibility, as a larger angle implies a larger domain of feasibility and can therefore be used to quantify the overall coexistence potential. It therefore provides a measure of the ease of coexistence for the whole community. Whenever the vector of intrinsic growth rates **r** fails to be within the domain of feasibility, at least one species goes extinct. This community-scale metric of coexistence Ω can be complemented by a species-scale measure *η* that captures the **resistance** of each species to disturbances. The directed angle between **r** and the two borders of the feasibility domain, denoted by *η*_1_ and *η*_2_, indeed quantifies the resistance of corresponding species toward extinction, as smaller angles indicate that the corresponding species is close to exclusion and that any perturbation of parameters would lead to its loss. These angles are set with a positive sign when **r** is inside the domain of feasibility, and the resistance measure goes to zero when the corresponding species goes toward extinction. In the example of figure 1A, species 2 is much closer to extinction than species 1.

The overall coexistence potential and resistance angles for two species can be computed as

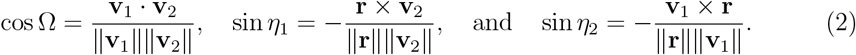

Note that while figure 1 is based on competition, this geometric approach can be used to depict any type of ecological interaction. Full details are provided in the Supporting Information S1, while figures S2, S3, S4, and S5 show other interaction types.

### 2.2 Evolution

Now that we have represented conditions of coexistence using geometrical methods, we aim at depicting the effects of eco-evolutionary dynamics using graphical tools in the same space (*r*_1_ − *r*_2_). The superposition of ecological and evolutionary components would then allow us to display how eco-evolutionary dynamics affect coexistence, as in figure 1B-C. To model eco-evolutionary dynamics, we rely on the adaptive dynamics framework (Dieckmann and Law, 1996; Geritz et al., 1998). While this framework focuses on phenotypic effects of evolution, therefore simplifying genetic aspects, it offers the advantage of having a complete description of fitness, grounded in ecological dynamics, including both frequency and density dependence (Dieckmann and Ferrière, 2004). This is especially critical when looking at the ecological implications of evolution, as we do here.

In adaptive dynamics, fitness is defined as the per capita growth rate of a rare mutant of phenotype *x*_*m*_ in an environment defined by the equilibrium of the resident population *x* (Metz et al., 1992). *Here, we do not aim at following the whole dynamics of a given phenotype, but rather to understand the direction of natural selection at a given point in time and how it likely affects coexistence for future generations of the system at hand. We therefore assume that the evolving species is species 1 and that the mutation alters a phenotype directly linked to a given interaction (e*.*g. α*_11_ or *α*_12_) (see also Loeuille (2010*b*)). Such a mutation can be linked to different kinds of trade-offs. We assume that the mutation either incurs allocative costs, in which case *r*_1_ is directly affected, or ecological costs (sensu Strauss et al. (2002)) in which case a mutation offering a benefit on an interaction will lead to costs on another. For instance, in figure 1B-C, we consider a mutation along the *r* − *K* continuum (MacArthur and Wilson, 1967; Pianka, 1970), in which case the intrinsic competitive ability of the species *α*_11_ is traded against intrinsic fecundity *r*_1_, which corresponds in our case to an allocative trade-off. Trade-offs may also be temporally absent (e.g. a trait has a cost in terms of specialist enemies that are not present, or an allocative cost in a context where much energy is available). In such instances, our method still applies, and evolution is simply directional.

Figure 1B shows such an example in which traits that decrease intraspecific competition are likely selected. As *α*_11_ becomes less negative, this affects the position of the vector **v**_1_ (blue arrow attached to the vector **v**_1_) with an allocative cost on the intrinsic growth rate *r*_1_ (red arrow attached to the vector **r**). To see why the geometry of this panel corresponds to a likely selection of competitive ability, let us compute the fitness gradient associated with such as scenario. Remembering that fitness is defined by the *per capita* growth rate of a rare mutant of trait *α*^*m*^ in a resident population of trait *α*_11_ (Metz et al., 1992), the fitness gradient, which gives the direction of evolution (Dieckmann and Law, 1996), can be written:

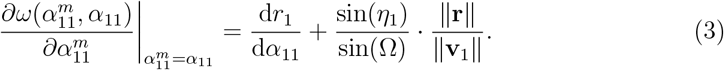

Therefore, K-strategies are favoured (higher *α*_11_ being selected when this gradient is positive, i.e. if the trade-off strength 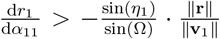, and counter-selected for the opposite inequality. Intriguingly, this means that adaptive dynamics can also be partly captured by the graphical approach. The ratio of angles and vector length 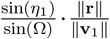, i.e., geometry of coexistence, indeed determine a threshold in the allocative cost for selection or counter-selection. For instance, in figure 1B, the angle *η*_1_ being quite large and the vector **r** having a large norm compared to the vector **v**_1_, selection of *K*-strategies is likely, as the contrary would require extremely high costs. Conversely, figure 1C shows an example where selection of *r*-strategies is expected, the angle *η*_1_ being quite small and the vector **v**_1_ having a large norm compared to the vector **r**. Full derivation is provided in Supporting Information S2.

In the case of increases in *α*_11_ (figure 1B), evolution harms coexistence both when measured based on overall coexistence potentials (Ω) and when measured as resistance of species 1 (*η*_1_). Not surprisingly, the opposite applies in the case of decreased *α*_11_ (figure 1C). Effects of evolution on species 2 resistance (*η*_2_) is undetermined, as it depends on the relative magnitude of the two blue and red arrows, as well as on the amplitude of vectors **r** and **v**_1_. Nevertheless, we can further compute its derivative relative to the intraspecific competition, again based on the geometry of coexistence and the trade-off strength,

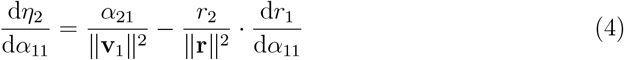

This expression gives us a second threshold in the trade-off strength, 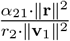 determining the direction of the change in species 2 resistance. Full derivation is provided in Supporting Information S2. Note that effects of evolution on *η*_2_ is undetermined because we are here in a competition context. If considering an interaction with a positive effect of species 1 on species 2, *α*_21_ *>* 0, and a positive intrinsic growth rate for species 2, *r*_2_ *>* 0, based on the previous equation, the direction of *η*_2_ evolution is systematically determined (see figure S3 panels B and D).

Our graphical approach, here shown for competitive interactions and allocative trade-offs, can be extended to any types of pairwise interactions (competition, antagonistic, and mutualistic), and trade-off types (allocative and ecological trade-offs), or even no trade-off. In Supporting Information S2, the reader may find the full case when the phenotype affects *α*_11_ and *r*_1_, in S3 the phenotype affects *α*_12_ and *r*_1_, in S4 the phenotype affects *α*_11_ and *α*_12_, and finally in S5 the phenotype affects *r*_1_, *α*_12_, and *α*_21_.

## 3 Applying the graphical approach to empirically informed cases

While the geometrical approach we propose can be used to illustrate and understand various theoretical cases (as in the previous part), it is also possible to apply it to various situations in which empirical information is known. Once the ecological stage is set (interaction type, number of species at hand), it is indeed possible to report known variations of key phenotypes and/or known or assumed trade-off constraints directly on the graphical approach to highlight possible implications of evolution for coexistence. We here highlight this for two types of traits that are particularly well-documented: plant defences and body size.

### 3.1 Evolution of plant defences and the control of invasive species

Our first illustrative example is based on the ideas expressed in Müller-Schärer et al. (2004) that investigate the evolution of defences of invasive plant species in their new range. The article proposes to distinguish two types of defence traits. Based on empirical evidence, *qualitative defences* (toxins) are assumed to be efficient against generalist enemies, but at the cost of attracting specialists. Specialists indeed use such defences as attraction cues, are not affected by these defences due to their tight co-evolution with the plant species, and may even use these toxins after ingestion as their own defences. The selection of qualitative defences therefore follows ecological trade-offs, balancing interactions with specialist vs generalist herbivores. Because specialists are often absent in the new range, qualitative defences become cost-free and higher levels can be selected there. *Quantitative defences* (e.g. tannins, lignin) are supposed to involve direct costs in terms of growth rate (Herms and Mattson, 1992) and/or competitive ability (Müller-Schärer et al., 2004), therefore involving both allocative costs and ecological costs. If, as stated above, qualitative defences have been selected, herbivore pressure should be low on the invasive species, and quantitative defences should be unnecessary and counter-selected (Müller-Schärer et al., 2004). This counter-selection would result in higher growth rates and higher competitive ability for the evolving invasive species.

While this empirical case may seem a bit complex (two different types of defences, with multiple impacts on ecological interactions and growth rates), our graphical approach can accommodate this type of complexity. To show this, we now plug this empirically-based analysis into our graphical approach. For clarity, we separate the effects of evolution into two parts: (i) on the invasive plant – generalist herbivore system (figure 2A-B) and (ii) on the invasive plant – local competitors (figure 2C-D). The graphical analysis highlights the crucial role of certain aspects of the problem in analysing the consequences of evolution for the maintenance of the system. More specifically, these consequences depend on the relative strength of ecological and allocative costs of defences (left vs right column of figure 2).

**Figure 2:**
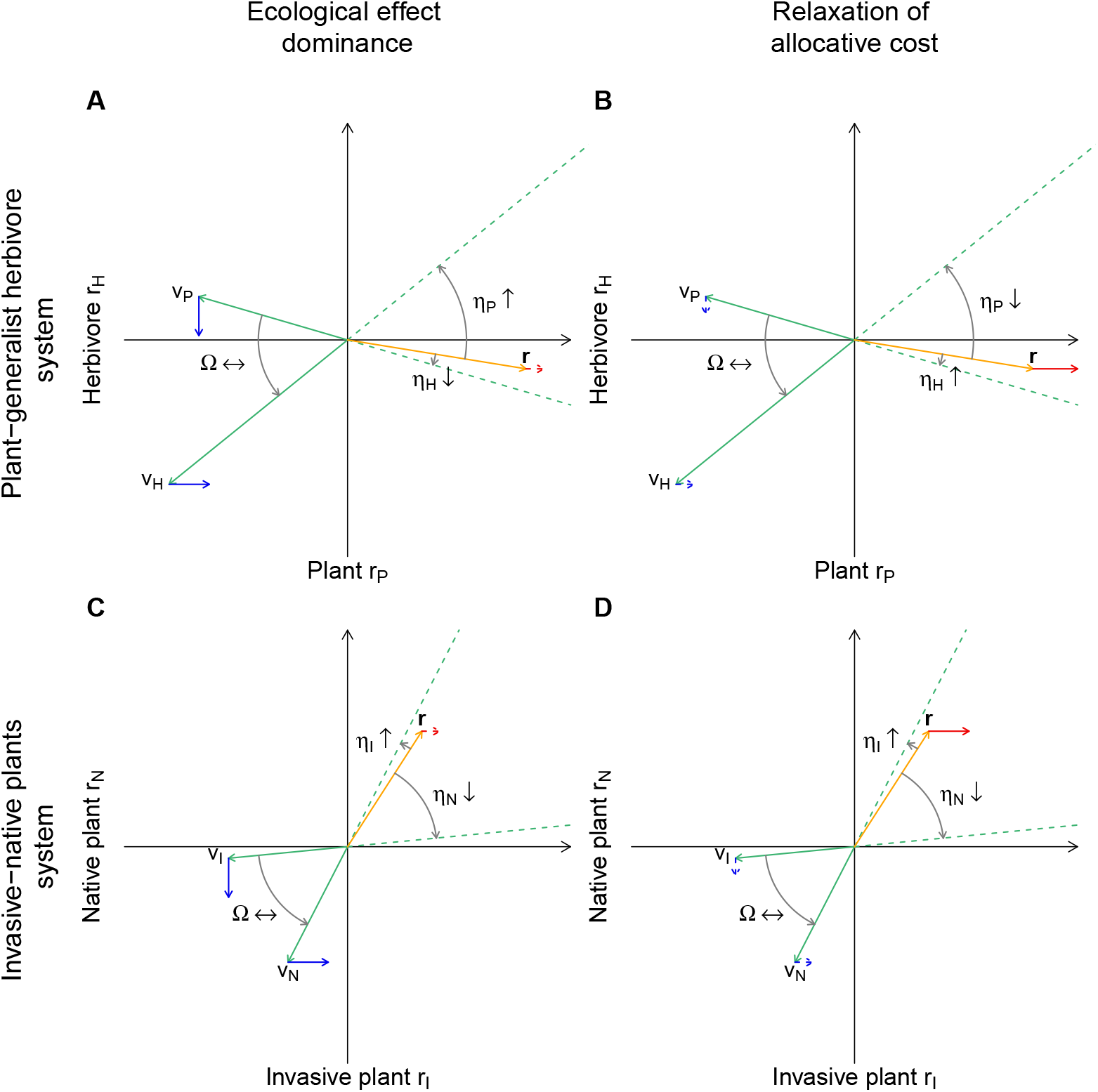
Effects of evolution of plant defences, based on empirical arguments proposed by (Müller-Schärer et al., 2004). Here, evolution is considered on two different types of defence traits: qualitative and quantitative, within the new range of the invasive species. Panels **A** and **B** show effects on coexistence from the herbivore–invasive plant perspective, while panels **C** and **D** show the effects of evolution from a native–invasive perspective. Strength of ecological vs allocation costs are contrasted, the former being dominant in **A** and **C**, the latter in **B** and **D**. Note that effects on coexistence depend on the relative costs in the plant–herbivore perspective, not in the invasive–native perspective.

First, consider the effects of the evolution of defences on the (invasive) plant-herbivore system. The evolution of toxins, based on allocation costs, decreases the impact of generalist herbivores on plants, thereby leading to the movements of vectors **v**_*P*_ and **v**_*H*_ (blue arrows on figure 2A-B). Because quantitative defences are counter-selected, the relaxation of associated allocation costs leads to higher plant growth rates (red arrow on figure 2A-B). Based on the ecological effects alone (blue arrows), note that we expect that evolution does not change much the overall coexistence at the community scale (as measured by Ω). However, consequences for herbivore and plant compartments vastly differ depending on the relative size of ecological and allocation costs. When ecological effects are dominant (figure 2A), evolution mostly harms the herbivore population (lower resistance *η*_*H*_) while enhancing plant robustness (higher *η*_*P*_). In such situations, direct effects of the evolution of toxins dominate; the trophic interaction is smaller, which leads to a reduction in energy transfer detrimental to the herbivore population. Conversely, if the dominant effect is the relaxation of allocative costs (i.e. higher *r*_*P*_, as in figure 2B), evolution fosters herbivore resistance (increase in *η*_*H*_) and decreases plant resistance (lower *η*_*P*_). Indeed, in such a situation, the counter-selection of quantitative defences allows an increase in plant basic productivity that increases the herbivore compartment through bottom-up effects.

When looking at the consequences of the evolution of invasive-native competition, ecological vs allocative costs now influence the outcome in synergy (figure 2C-D). Remembering that the counter-selection of quantitative defences not only relaxes allocation costs (red arrow on figure 2C-D), but also increases the competitive ability of the invasive species (blue arrows on figure 2C-D), consequences for the system persistence can be directly analysed using the graphical approach. When ecological costs are dominant (figure 2C), evolution leads to a decrease in native resistance (lower *η*_*N*_) and to an increase in invasive resistance (higher *η*_*I*_), as expected, based on the increased competitive asymmetry. This effect is reinforced by the relaxation of allocative costs in figure 2D. The invasive plant then cumulates two advantages: higher competitive ability and higher growth rates, further reducing native resistance and threatening the maintenance of the system.

Our graphical approach is used here to provide possible scenarios for future coexistence. However, it also serves to point out critical aspects that should be investigated to get a more in-depth understanding of the evolution and maintenance of these systems. Here, for instance, the outcome on the trophic side (plant-herbivore) depends on the relative size of ecological and allocative costs, while the competition outcome is qualitatively the same (though increased when allocative costs are important). While the assessment of trade-offs is notoriously difficult, note that quite a lot of information exists on both types of costs (e.g. Herms and Mattson (1992); Müller-Schärer et al. (2004)), so that informed scenarios can be proposed when this information is available. If not, our approach highlights that getting information on this relative size of costs could be a priority. Furthermore, while figure 2 proposes that Ω may be left unchanged throughout evolution, this guess is based on that we have no information on the asymmetry of trait impacts on the interactions (i.e. we assumed similar sizes for blue arrows). Getting further information on such impacts would allow us to refine possible consequences of evolution.

### 3.2 Ecological effects of body size evolution

Body size (most often measured as adult body mass) is likely the best documented phenotype in ecology. Variations in body size are known to affect many aspects of life history, metabolic constraints and ecological interactions (Peters, 1986). Based on all these empirical observations, we propose that it is possible to use the graphical approach to better understand how the evolution of body size influences coexistence in ecological communities. To do so, we build on two examples: (i) the often reported reduction in body size due to intense fishing (e.g. in cod populations (Olsen et al., 2004)), (ii) the general idea that larger body size offers a competitive advantage in interference competition (Persson, 1985). These two examples are chosen because they allow us to illustrate different types of interactions (trophic and competitive respectively) as well as a phenotype that incurs various benefits and costs (hence complex trade-off shapes).

Olsen et al. (2004) suggests a fast evolution of cod populations in Newfoundland due to intense fishing. Given the high rate of mortality due to direct exploitation, individuals that mature early have a selective advantage. This leads to a fast reduction of the age at maturity, accompanied by a reduction in adult body size. Such an evolution is not an exception, and has been observed in various harvested or hunted populations (Barot et al., 2004; Grift et al., 2003; Conover et al., 2005). Our graphical approach allows the investigation of the possible consequences of the evolved reduction in cod body size (Olsen et al., 2004) for coexistence with their prey. Following the metabolic theory of ecology (Brown et al., 2004), we assume *r*_*P*_ is decreased (i.e. faster decline) when body size becomes smaller. We also assume that trophic interactions are based on relative sizes of predators and prey (Brose et al., 2006; Naisbit et al., 2012; Li et al., 2023). Assuming that the cod population is initially adapted to the size of its prey, evolved reductions in body size come at a cost in terms of attack rate.

These assumptions lead to the graphical outcome reported on figure 3A. Since we assumed that the trophic interaction is here relaxed due to changes in predator body size, the impact on interaction vectors is arguably asymmetric. Indeed, only a fraction of consumed prey is incorporated in predator variations (Lindeman, 1942), making predator vs prey impacts intrinsically asymmetric (Neutel and Thorne, 2014). We hence assume that impacts on vector **v**_*P*_ is larger than impacts on vector **v**_*N*_. Under this hypothesis, overall coexistence is enhanced by evolution (as measured by angle Ω). Effects on the resistance of the two trophic levels depend on whether changes in body size affect more strongly the basic growth rate of the predator (red arrow) or the trophic interaction (blue arrows). In the first case, predator resistance (*η*_*P*_) is increased while prey resistance *η*_*N*_ decreases. Higher predator growth rate indeed increases top-down pressure on prey population, and they benefit from lower mortality. In the second case, as displayed on panel A, predator resistance is decreased while prey resistance is increased. Due to fishing pressures, the size of the predator is increasingly maladapted to the consumption of the prey species and top-down control is reduced.

**Figure 3:**
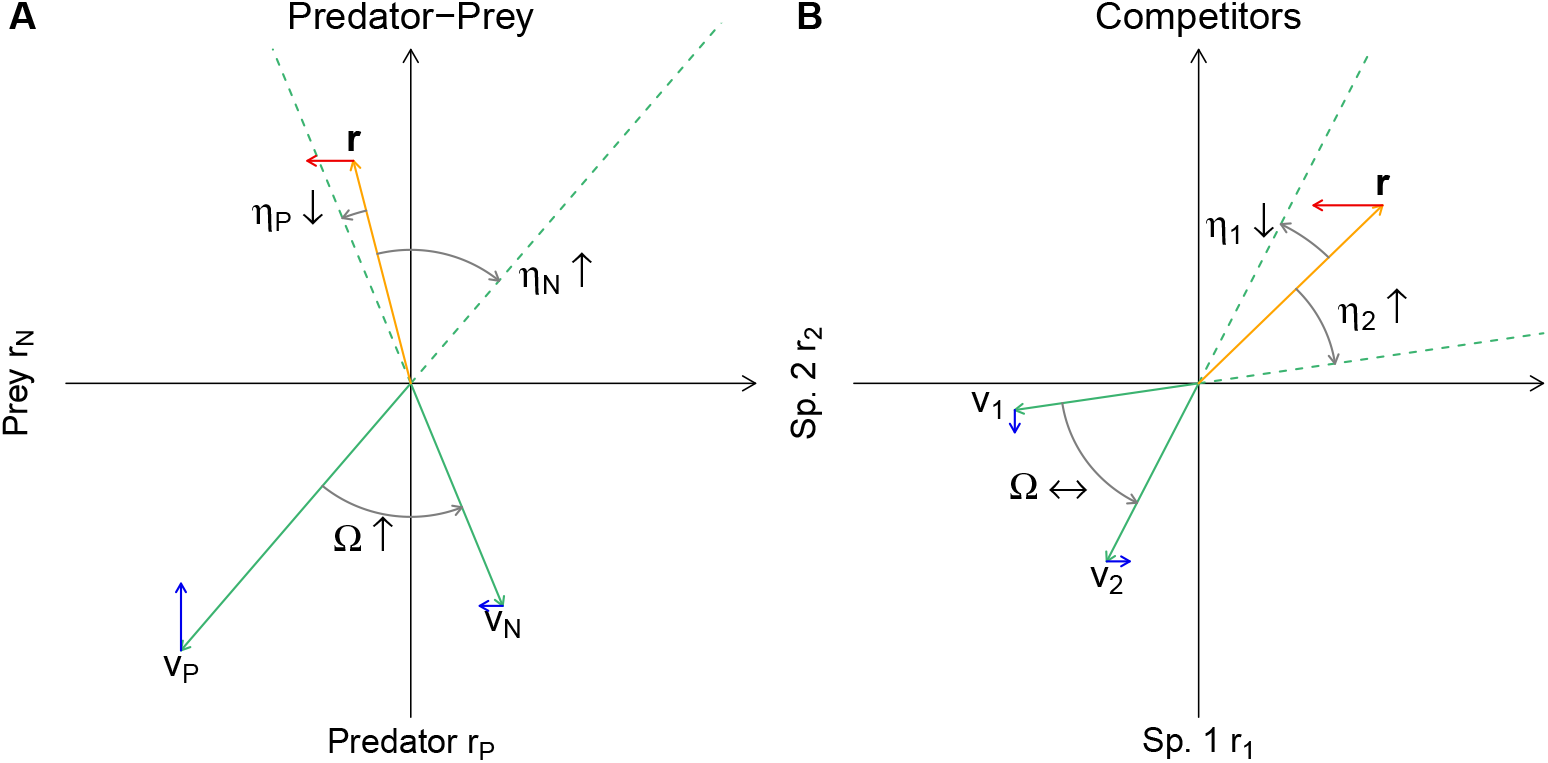
Effects of the evolution of body size (as measured by adult body mass) on species coexistence. Panel **A** trophic. Based on the evolution of lowered body size in harvested fishes. E.g. reduction in cod (predator) body size based on (Olsen et al., 2004). We here assume that such an evolution decreases the cod attack rate (blue arrows on interaction vectors), while also modifying Malthusian growth rates (based on metabolic theory, red arrow). Evolution here enhances overall coexistence, while making the predator more vulnerable. Panel **B** competition. We assume intense interference competition, selecting for higher body sizes. Such an evolution modifies the competitive hierarchy (blue arrows on interaction vectors) while reductions of the Malthusian growth rate are again based on the metabolic theory of ecology (red arrow). Note that we here assume that interaction impacts are low compared to growth variations, so that evolution increases the evolving species vulnerability, while leaving overall coexistence potential unchanged.

Effects of variations in body size in a competitive context can be similarly worked out. Imagine that competition is asymmetric, large body sizes being favoured (Persson, 1985). For instance, in Trinidadian guppy populations, a long-term study on four different populations highlighted how larger body size is systematically linked to increased competition on smaller phenotypes (Griffiths et al., 2020). Assuming that resources are limited, competition intense, so that such larger body sizes are selected for species 1, it is possible to infer the consequences of such variations using our framework. More precisely, still based on the metabolic theory of ecology (Brown et al., 2004), we assume that such an increase in body comes at the cost of a lower intrinsic growth rate. As shown on figure 3B, evolution is leading to a shift of vectors **v**_1_ and **v**_2_, so that overall coexistence (angle Ω) is not changed in a systematic way. Concerning species resistance, as in the previous case, the outcome depends on the relative effects of variations of body size on the demographic (red arrow) vs competitive interactions (blue arrows). If demographic components vary more with body size, as displayed on 3B, evolution may erode the existence of the evolving species (evolutionary deterioration *sensu* Matsuda and Abrams (1994); Dieckmann and Ferrière (2004)) while if movement of vector **v**_2_ (linked to the fact that competitive effects of species 2 on species 1 are now relaxed) is important, resistance of species 1 is higher given the observed variations in body size.

## 4 Discussion

Understanding the intricate relationships between evolution and the diversity and structure of ecological communities is at the heart of eco-evolutionary studies (Edwards et al., 2018; Hart et al., 2019). Given the many observations of evolution recently reported under current changes (Bonnet et al., 2022), this understanding is urgently needed to better forecast the future of biodiversity and its implications for ecosystem functioning. While the method we propose here is based on just a few species, we believe that it may help by providing a graphical approach that explicitly links (i) evolutionary dynamics, here analysed through an adaptive dynamics approach; (ii) coexistence theory, based on structural stability. While mathematical analysis incorporating these two frameworks is certainly possible (e.g. Lepori et al. (2024)), our graphical approach offers the possibility to be readily usable by non-theoreticians, thereby possibly facilitating the discussion between experimental/empirical observations and theoretical developments of eco-evolutionary dynamics.

Concerning the evolutionary part, we show that the fitness gradient can be decomposed in two parts: one that captures the costs of the phenotypic trait at hand (for instance, how the intrinsic growth rate changes when the trait varies (allocation costs)) and the other that can be easily expressed in terms of basic growth rates and interaction intensity. This second part can be directly drawn within our graphical approach, that is grounded in the representation of interaction vectors within a space defined by the growth rates of species. Even though assessing the amplitude of evolutionary costs is notoriously difficult (Mauricio, 1998; Reznick et al., 2000), the graphical approach may still give the likely direction of evolution. For instance, figure 1 (panels B and C) shows two situations in which selection and counter-selection of the traits are likely, in the sense that only extreme costs could prevent these directions. We, however, emphasize that in between these two extreme situations, the determination of the likely direction of evolution may be more ambiguous. We also stress that while we here developed our evolutionary approach based on adaptive dynamics approaches, it is conceivable to use it under other frameworks. For instance, in many quantitative genetic studies, fitness is expressed based on variations of ecological characteristics of the species and of its interactions (e.g. Taper and Case (1992); Norberg et al. (2012). In such conditions, the fitness gradient may similarly be computed, and the approach we propose applied.

Once the direction of evolution is found out, it can be displayed directly within the graphical approach, as arrows affecting the three vectors that determine coexistence. The recent development of structural stability analysis (Rohr et al., 2014; Saavedra et al., 2017; Lepori et al., 2024) can then be used to determine how the general coexistence (Ω angle) and the vulnerabilities of the two species are changed through evolution. In some cases, the role of evolution is not ambiguous. For instance, the example based on evolution of defences in invasive plants show that vulnerabilities change in a systematic way, both for the invasive and for the resident species (figure 2). The same case, however, suggests that general coexistence may not be affected (angle Ω is mostly unchanged). We acknowledge however that this statement is based on the idea that evolution affects the two interaction vectors roughly the same way. To provide a more precise assessment, one would need to uncover the relative amplitude of evolutionary effects on the various interaction rates, which may be empirically difficult. Still, the graphical approach proposes a reference scenario, that can then be tested experimentally or through an analysis of empirical datasets.

The method can only be applied based on the assessment of interaction rates and of intrinsic growth rates of species. Information may however be limiting, especially regarding the intensity of ecological interactions. While the knowledge of ecological interaction intensity is sparse, many methods have been developed throughout the years, from experimental approaches dating back to Paine (1992) to the reconstruction of the variability of feeding links within food webs (Paine, 1980; Rooney et al., 2006; Goldwasser and Roughgarden, 1993). Methods based on steady state conditions and species allometry (including ecosim/ecopath approaches) are vastly used to assess relative interactions (De Ruiter et al., 1995; Jacquet et al., 2016) (see Gauzens et al. (2023) for a recent package development). When time series of abundances are available, more advanced techniques can also be used (Sugihara, 1994; Nguyen et al., 2025). If a precise assessment of ecological interactions cannot be achieved, the method may still allow discussing plausible scenarios and offer a first take on the question at hand. For instance, in the various empirical examples we propose in the text, a precise assessment of the interaction is not included, but we simply use available information on reported evolutions to propose possible effects on coexistence. More precise assessment of the various ingredients would, however, allow us to refine the proposed outcomes.

The framework we propose has several important limitations. First, as a graphical approach, it is suited to study subsystems containing few species. Typically, the coexistence space is easy to draw in two dimensions (all figures presented here) and could be extended to a third dimension to depict three species at the same time. For more complex systems (diffuse co-evolution *sensu* Janzen (1980)), a graphical approach may not be suitable. A possible way around is to split the complex system in different subparts and to try to understand various pieces of the puzzle. For instance, in the invasiveresident-herbivore system we use here, based on empirical inputs from Müller-Schärer et al. (2004), we split the system in two two-species subsystem, to allow a better visualization of the effects of evolution within guilds and among trophic levels. For higher diversity scenarios, however, numerical simulations are likely needed, and the graphical approach becomes limited. We also restate that the method is used to tackle effects and direction of evolution at a given point in time, not to follow the evolutionary dynamics until their outcome. While this limit is quite restrictive from a purely theoretical perspective (e.g., understanding the eventual effects of evolution, close to the evolutionary attractor), the framework may still help to work out the effects of evolution on coexistence, based on experiments or empirical data, for the next generations of the considered species. In turn, this information may be used and considered for a better understanding and management of the systems at hand.

A development of proper graphical tools is often instrumental to improve the link between theory and its applications. Lotka-Volterra systems of competition and predation, while often mathematically tractable, are often taught based on isocline graphs, and these isocline graphs allow an intuition that serves to make theory accessible to a broad spectrum ranging from bachelor students to non-theoretician researchers in ecology. Even among theoreticians, simplified graphical approaches help to clarify key mechanisms, as exemplified by the use of ZNGI (zero net growth isoclines) approaches throughout decades (Tilman, 1982; Koffel et al., 2021). On the evolution theory side, adaptive dynamics is a good example of a modelling technique that succeeded not only based on its theoretical merits (e.g. an ecologically based definition of fitness Metz et al. (1992) and of the related variations of traits Dieckmann and Law (1996)), but also because a graphical approach was proposed to visualize and understand evolutionary dynamics based on Pairwise Invasibility Plots (Geritz et al., 1998). We therefore believe that a graphical approach that link evolution and coexistence may help to bridge a gap between theoretical developments and empirical assessments of the question, and hope that the framework we propose here will be useful to fellow researchers interested in these questions.

## Acknowledgments

RPR acknowledges the Swiss National Science Foundation Sinergia grant no CRSII5 202290, and Swiss National Science Foundation — U.S. National Science Foundation, lead-agency grant no 320030L-227556

## S1 Geometry of two species coexistence

We consider the Lotka-Volterra model for two interacting species of abundance *N*_1_ and *N*_2_ (Volterra, 1931; Case, 2000). It is given by the following set of differential equations:

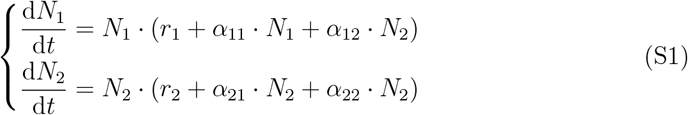

The parameters are the intrinsic growth rates *r*_*i*_ of species *i*, and the *per capita* interaction of species *j* on species *i* denoted by *α*_*ij*_. These parameters can be encapsulated into the vector **r** of intrinsic growth rates and the matrix **A** of *per capita* interactions:

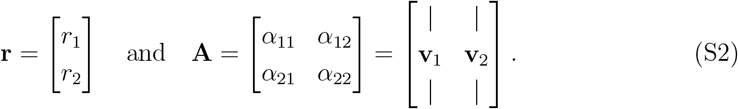

The interaction matrix can be decomposed into two column vectors **v**_1_ and **v**_2_ capturing the *per capita* effect of species 1 and 2 on their interactor and on themselves.

To allow coexistence in such a system, two conditions need to be fulfilled: (i) feasibility and (ii) stability (Volterra, 1931; MacArthur and Levins, 1967; Vandermeer, 1975; Logofet, 1993; Case, 2000). Feasibility corresponds to the existence of a nontrivial, i.e., positive, equilibrium point. Such a point denoted by 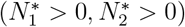 must, therefore, be a solution of the following linear equation.

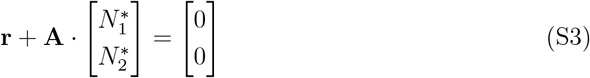

The stability conditions, in turn, grant that the non-trivial equilibrium, if it exists, is locally stable. The condition for stability is given by

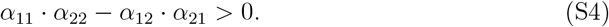

Note, that in a two species Lotka-Volterra model, local stability is equivalent to global stability.

The aim of the structural approach to coexistence is to provide a geometric interpretation to the feasibility (equ. S3) and stability (equ. S4) equations. Regarding the feasibility constraints, instead of solving directly the equation (S3), we represent the set of intrinsic growth rates (**r**) leading to a positive solution. This is achieved by rewriting equation (S3) as

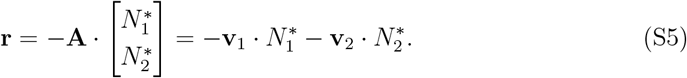

That is, as feasibility requires 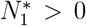 and 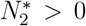 0, the set of **r** leading to a feasible solution are the positive linear combinations of the vectors −**v**_1_ and −**v**_2_. This set of feasible **r**, called the feasibility domain, can be represented geometrically as a cone (figure S1) in the two-dimensional *r*_1_ − *r*_2_ space, where the x-axis is the intrinsic growth rate of species 1 and the y-axis is the intrinsic growth rate of species 2. The borders of the feasibility domain are determined by the *per capita* interaction strengths. The smallest angle between the two vectors **v**_1_ and **v**_2_, denoted by Ω, quantifies the size of the feasibility domain and therefore the potential of a feasibility.

**Figure S1:**
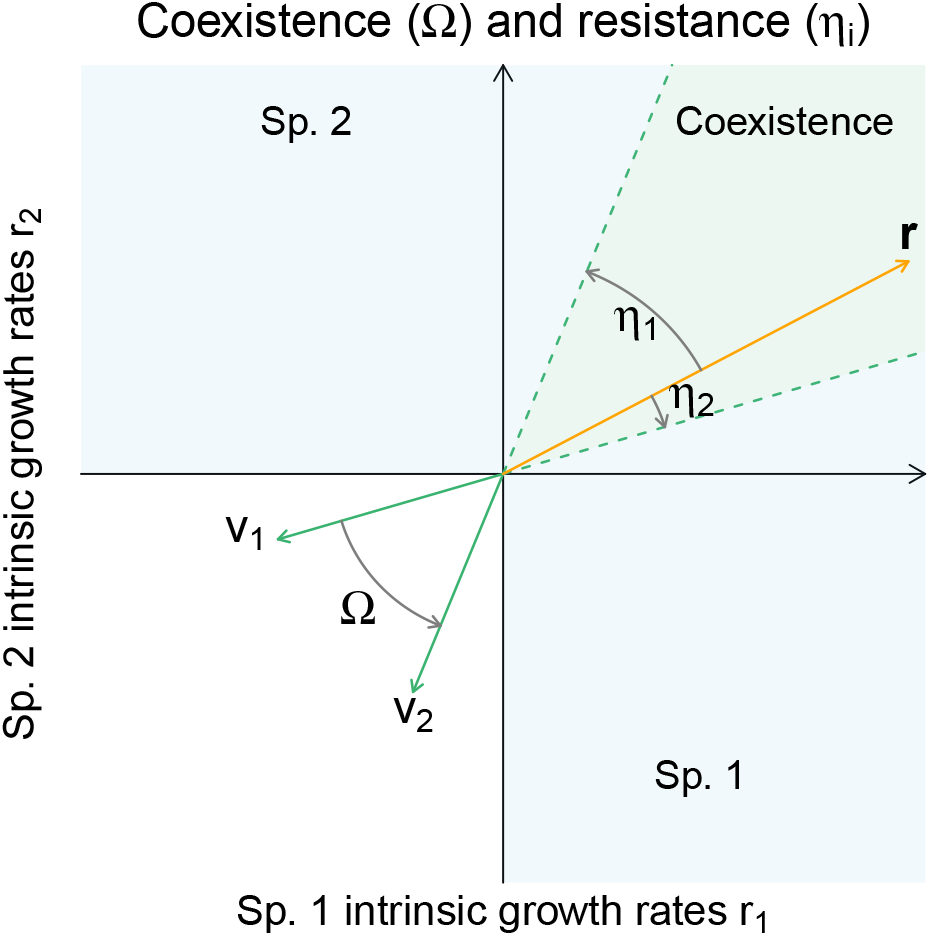
Geometric approach to coexistence. The per capita interaction strength determine the vectors **v**_**1**_ and **v**_2_, while, **r** gives the intrinsic growth rates. The green cone is the domain of feasibility, i.e., feasibility is granted if and only if **r** is inside this domain. The directed angles Ω determine the overall coexistence potential and stability condition, while the angles *η*_1_ and *η*_2_ are the resistance angles of species 1 and 2, respectively.

In turn, the stability condition also has a geometric representation. The term *α*_11_ · *α*_22_ − *α*_12_ · *α*_21_ in the stability condition, is equivalent to the cross product between the two vectors **v**_1_ and **v**_2_, and therefore, proportional to the sinus of Ω:

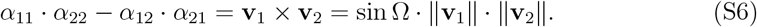

That is, stability is equivalent to a positive cross product between the vectors **v**_1_ and **v**_2_, while a negative cross product implies instability. Therefore, to geometrically represent stability, we can add a direction to the angle Ω, i.e., turn it into a directed angle between **v**_1_ and **v**_2_. The direction of Ω is given in figure S1 by the arrow on the arc of a circle depicting Ω. The directed angle Ω is defined as the smallest angle between **v**_1_ and **v**_2_, and its sign is positive if the vector **v**_1_ is before the vector **v**_2_ when considering a counter-clockwise direction. Otherwise, the sign is negative. Consequently, stability is granted if the directed angle is positive but smaller than 180^°^, i.e. 0^°^ *<* Ω *<* 180^°^.

Assuming that stability conditions are fulfilled, if the vector of intrinsic growth rates **r** falls outside the feasibility domain (green cone), at least one of the two species goes extinct. In particular, if **r** is located in one of the two blue areas (figure S1), then only species 1 or species 2 survive, while the other goes extinct. Therefore, as represented in figure S1, the two directed angles *η*_1_ and *η*_2_ between **r** and the border of the feasibility domain are called species resistance. Note that the sign of the directed angle *η*_1_ is defined counter-clockwise (like Ω), while the sign of the angle *η*_2_ is defined clockwise (opposite direction). This allows the interpretation that (1) the smaller the angle (but still positive), the closer the corresponding species is to extinction, and (2) feasibility is granted if and only if both resistance angles are positive, while if at least one angle is negative, the corresponding species is extinct.

The overall coexistence potential angle Ω and the two resistance angles *η*_1_ and *η*_2_ define the metrics of coexistence and are computed as follows:

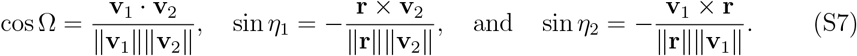

Under feasibility and stability conditions, the equilibrium densities can be computed from the overall coexistence potential and resistance angles, i.e., have a geometric perspective:

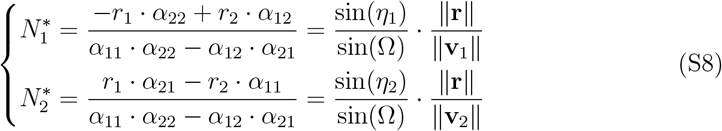

The middle terms of the above set of equations are the classical solution of equation (S3). Now, at the denominator, we can identify the term determining the stability condition, which is equivalent to the cross product between **v**_1_ and **v**_2_, i.e, proportional to the sinus of Ω. While, the terms at the two numerators are the same as the cross product between **v**_2_ and **r** and between **v**_1_ and **r**, respectively. That is, the two numerators are proportional to the resistance angles. All that together leads to the right terms of equation set (S8).

Finally, note that this geometric approach can be applied to any type of interaction, as demonstrated in figure S2. The *r*_1_ − *r*_2_ space can be decomposed into four orthants. We can always assume negative density dependence of a species on itself; that is, the intraspecific interaction can be assumed to be negative *α*_11_ *<* 0 and *α*_22_ *<* 0. This implies that the vector **v**_1_ can only be in orthant II and III, while **v**_2_ can only be in orthant III and IV. This leads to four combinations, as illustrated by panels S1.A to S1.D describing all types of pairwise interactions.

**Figure S2:**
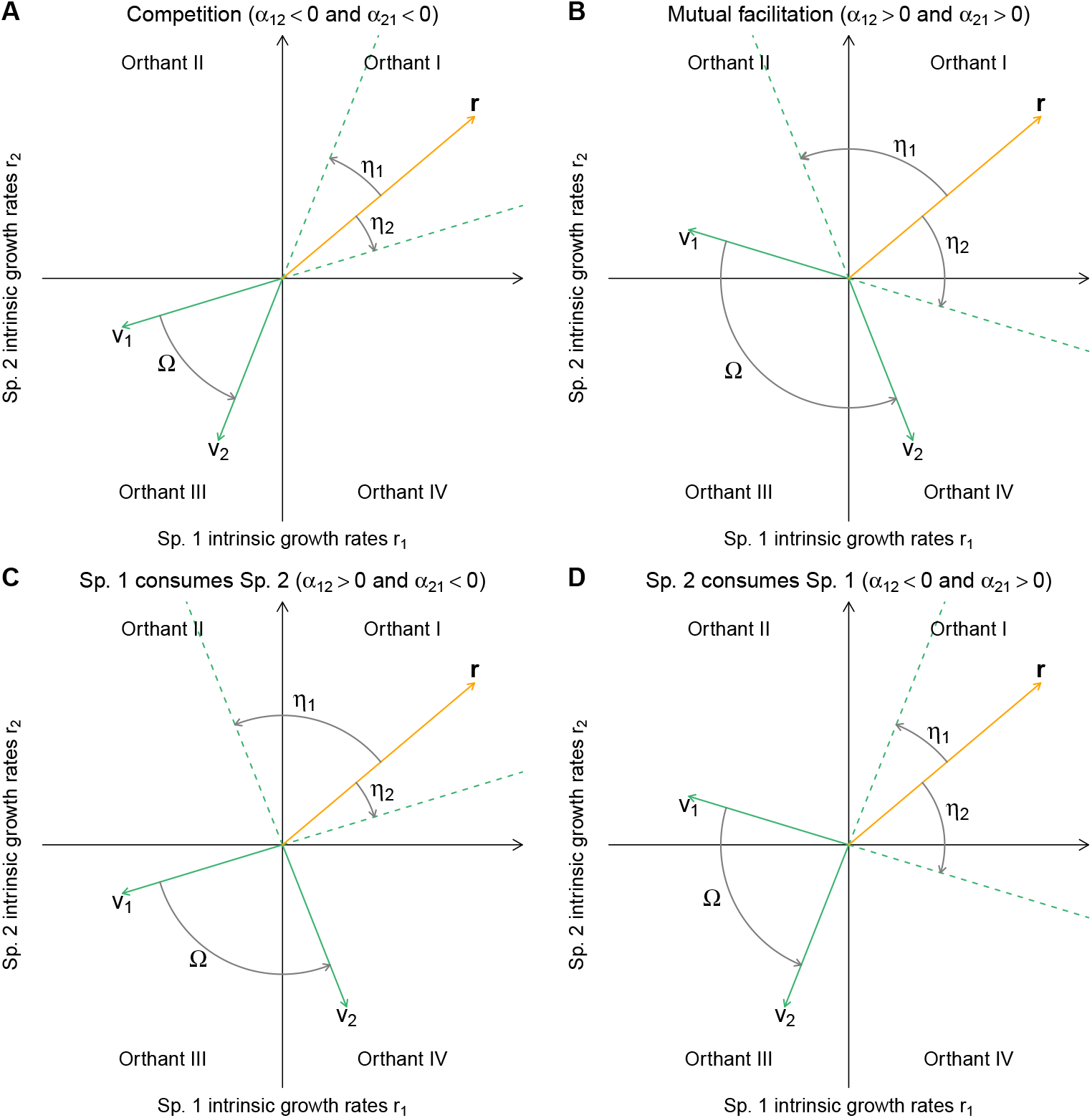
The position of the vectors **v**_1_ and **v**_2_ determines the types of interactions between species 1 and species 2.

## S2 Evolution of a phenotype affecting *α*_11_ and *r*_1_

Here, we assume that the trait *z* of species 1, which is under selection, impacts both the intrinsic growth rate *r*_1_(*z*) and intraspecific competition *α*_11_(*z*). Figure S3 shows the graphical analysis of the consequences of evolution on the geometry of coexistence for the four possible combinations of interactions. In this figure, we assume that evolution relaxes intraspecific competition at the cost of intrinsic growth rate, which relates, for example, to the classical *r* − *K* theory (MacArthur and Wilson, 1967). That is, evolution toward *K*-strategies results in a simultaneous decrease in *r*_1_ (red arrow) and increase in *α*_11_ (that therefore becomes less negative, blue arrow on vector **v**_1_). Other cases can be obtained simply be first choosing the appropriate direction of the red and blue arrows, and then deducing the direction of the black arrows. The figure shows, for this scenario, that the effects of evolution on overall coexistence potential Ω and on species 1 resistance *η*_1_ are systematically determined by the type of interaction that species 1 has with species 2. However, the effect on species 2 resistance *η*_2_ is more difficult to determine graphically, as both the vector **r** and **v**_1_ are displaced through evolution and, in some cases, in the same direction (panels A and C). In such instances, as mathematically demonstrated below, the impact of evolution on *η*_2_ is determined by the combination of the trade-off strength, d*r*_1_(*z*)/d*z /* d*α*_11_(*z*)/d*z*, and the vectors **r** and **v**_2_.

These graphical results can be mathematically proven by computing the derivative, relative to the trait *z*, of the overall coexistence potential Ω and the two resistance angles *η*_1_ and *η*_2_. Moreover, the selection gradient can also be related to the geometry of coexistence. We start by deriving the selection gradient, and then we compute dΩ/d*z*, d*η*_1_/d*z*, and d*η*_2_/d*z*.

The invasion fitness of a rare mutant of trait *z*_*m*_ in a resident population of trait *z* is given by its per capita growth rate when rare, the resident population being at equilibrium (Metz et al., 1992), which results in

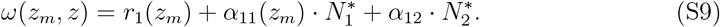

The direction of trait evolution is then given by the selection gradient, a positive selection gradient leading to the selection of higher trait values, while a negative gradient conversely leads to smaller trait values (Dieckmann and Law, 1996). Given the definition of fitness above, the selection gradient equals:

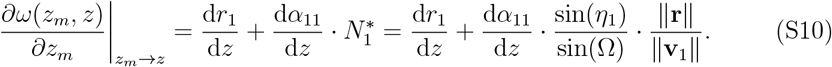

**Figure S3:**
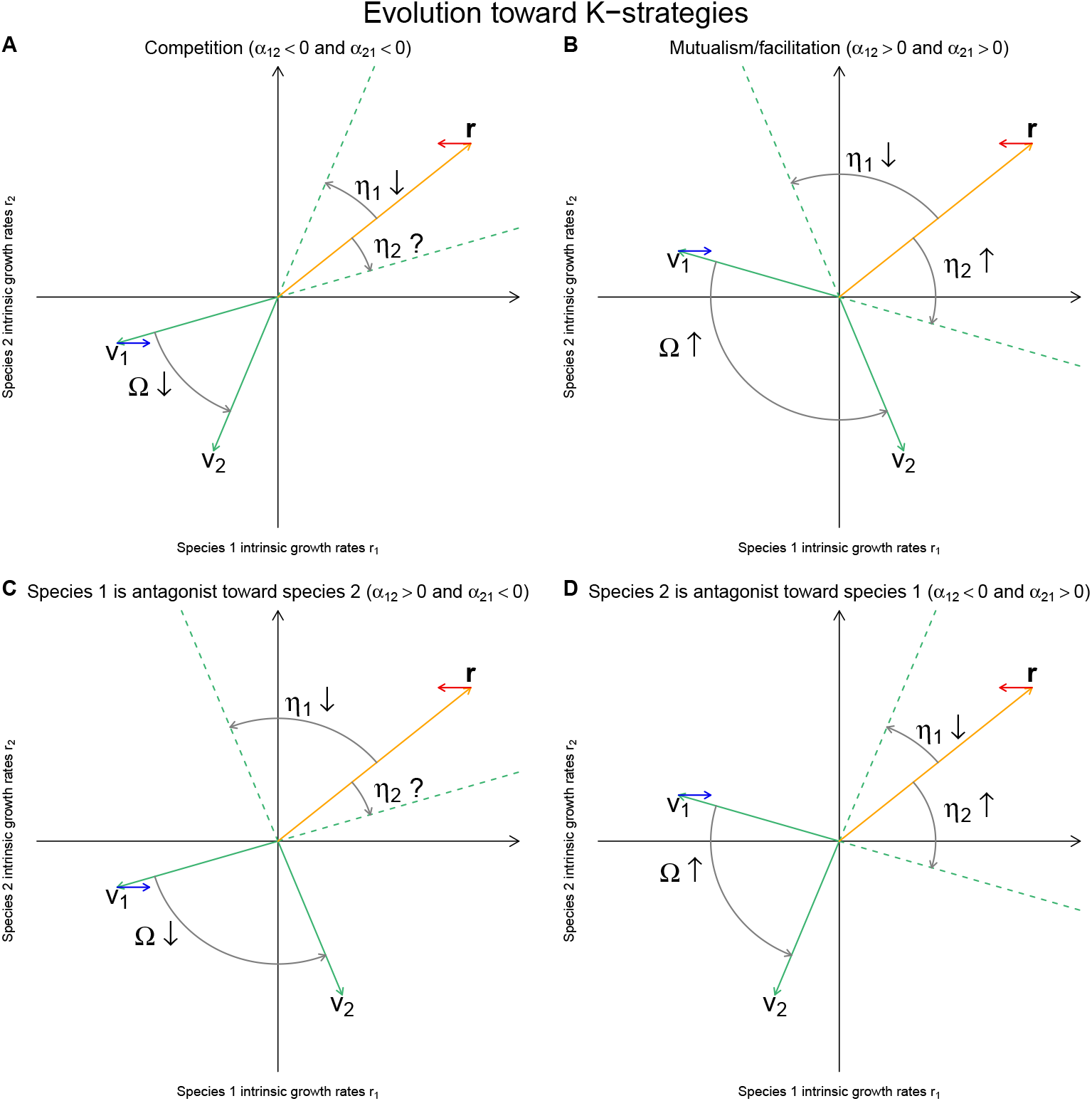
Effect of evolution of species 1 on the structural metrics in the case of an allocative trade-off between *r*_1_ and *α*_11_. Here, we assume that evolution relaxes intraspecific competition at the cost of a decrease in intrinsic growth rate (evolution toward a *K* strategy). Other cases can be obtained by simply setting accordingly the red and blue arrows, and then, deducing the direction of the black arrows.

Two cases for the direction of evolution occurs. First, one can assume an allocative tradeoff between *r*_1_(*z*) and *α*_11_, i.e., the sign of d*r*_1_(*z*)/d*z /* d*α*_11_(*z*)/d*z <* 0 is negative. In such case, the direction of evolution is determined the strength of this trade-off and the geometric of coexistence given by the angles Ω and *η*_1_, as well as the length of the two vectors, the vectors **r** and **v**_1_. In the second case, where no trade-off is assumed, the signs of d*r*_1_(*z*)/d*z* and d*α*_11_(*z*)/d*z* are the same, and therefore, the direction of evolution is determined by their sign.

The effects of evolution on the different metrics of coexistence are given by:

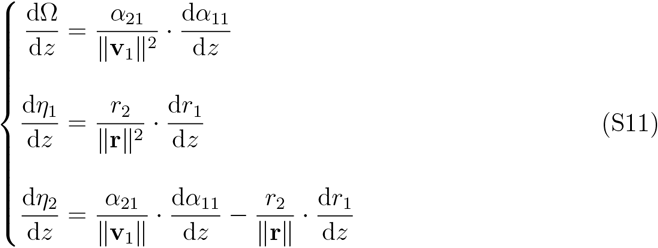

Note that dΩ/d*z* = d*η*_1_/d*z* + d*η*_2_/d*z*, which is coherent as, Ω = *η*_1_ + *η*_2_.

To easily relate these three equations to figure S3, we can rewrite them to make apparent the changes in *ω*(*z*_*m*_, *z*), Ω, *η*_1_, and *η*_2_ relative to the change in *α*_11_. This can also be considered equivalent to considering *z* = *α*_11_ and, consequently, *r*_1_ being a function of *α*_11_. This leads to

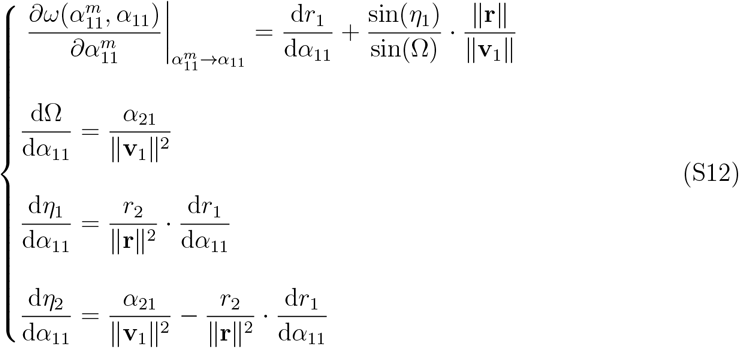

The first equation shows that *α*_11_ is selected if and only if sin(*η*_1_)*/* sin(Ω)·∥**r**∥*/*∥**v**_1_∥ *>* − d*r*_1_/d*α*_11_. That is, the geometry of coexistence (the second term of this equation) determines a threshold for selection and counter-selection in the trade-off case (i.e. when d*r*_1_/d*α*_11_ *>* 0). The second equation demonstrates that the evolution of *α*_11_ and Ω are in the same direction when the effect of species 1 on species 2 is positive (*α*_21_ *>* 0), as illustrated by panels B and D in figure S3, while it goes in the opposite direction when species 1 negatively affects species 2 (*α*_21_ *<* 0), as illustrated by panels A and C. The third equation demonstrates that whether the evolution of *α*_11_ and *η*_1_ are in the same or opposite direction is determined by the sign of *r*_2_ · d*r*_1_/d*α*_11_. As, on figure S3 we assumed d*r*_1_/d*α*_11_ *<* 0 and *r*_2_ *>* 0 evolution is in the opposite direction. The reader can imagine other cases, by, for instance locating the vector **r** in the fourth orthant on panels B and D. Finally, the last equation shows that the determination of the direction of evolution of *η*_2_ is more tricky, as already shown in figure S3, as it will depend on the signs of *r*_2_ *>* 0, *α*_21_ *>* 0, and d*r*_1_(*z*)/d*z /* d*α*_11_(*z*)/d*z*, as well as on the amplitude of the vectors **r** and **v**_1_. In some particulars cases, as in panels B and D, the direction is uniquely determined.

## S3 Evolution of a phenotype affecting *α*_12_ and *r*_1_

Here, we assume that the trait *z* of species 1, which is under selection, impacts both the intrinsic growth rate *r*_1_(*z*) and interspecific interaction *α*_12_(*z*). Figure S4 shows the graphical analysis of the consequences of evolution on the geometry of coexistence for the four possible combinations of interactions. In this figure, we assume that evolution either relaxes the negative effect of species 2 on species 1 (panels A and D), or increases the positive effect of species 2 on species 1 (panels B and D), both at the cost of intrinsic growth rate. Such a situation can, for instance, relate to when prey invest in defence, i.e., a reduction of the negative impact of the predator that translates to an increase in *α*_12_(*z*), at the cost of a decrease in its intrinsic growth rate *r*_1_(*z*) (e.g. for plant defences Herms and Mattson (1992)). That is, evolution results in a simultaneous decrease in *r*_1_ (red arrow) and increase in *α*_12_ (that therefore becomes less negative or more positive, blue arrow on vector **v**_2_). Other cases can be obtained simply be first choosing the appropriate direction of the red and blue arrows, and then deducing the direction of the black arrows. The figure shows, for this scenario, that the effects of evolution on overall coexistence potential Ω and on species 2 resistance *η*_2_ are systematically determined. However, the effect on species 1 resistance *η*_1_ is more difficult to determine graphically, as both vectors **r** and **v**_2_ are displaced through evolution and in the same direction. In such instances, as mathematically demonstrated below, the impact of evolution on *η*_1_ is determined by the combination of the trade-off strength, d*r*_1_(*z*)/d*z /* d*α*_12_(*z*)/d*z*, and by the norms of vectors **r** and **v**_2_.

These graphical results can be mathematically proven by computing the derivative, relative to the trait *z*, of the overall coexistence potential Ω and the two resistance angles *η*_1_ and *η*_2_. Moreover, the selection gradient can also be related to the geometry of coexistence. We start by deriving the selection gradient, and then we compute before computing the dΩ/d*z*, d*η*_1_/d*z*, and d*η*_2_/d*z*.

Following the same rational as in section S2, the invasion fitness of a rare mutant of trait *z*_*m*_ in a resident population of trait *z* is given by

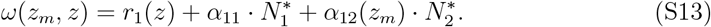

The direction of trait evolution is then given by the selection gradient

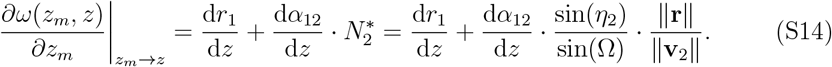

**Figure S4:**
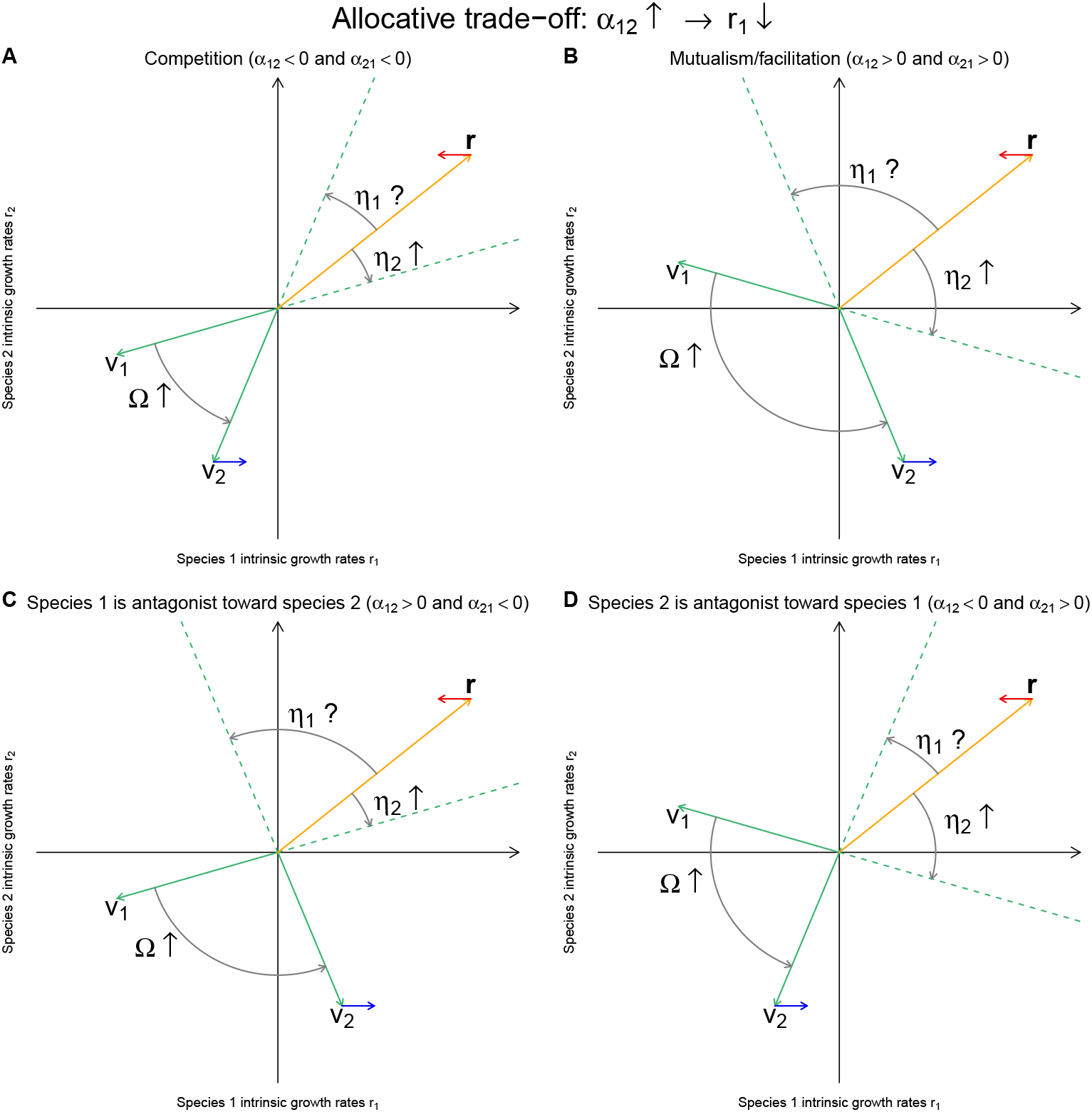
Effect of evolution of species 1 on the structural metrics in the case of an allocative trade-off between *r*_1_ and *α*_12_. Here, we assume that evolution relaxes the negative effect of species 2 on species 1 (evolution toward better defences), or increase the positive effect, at the cost of a decrease in intrinsic growth rate. Other cases can be obtained by simply setting accordingly the red and blue arrows, and then, deducing the direction of the black arrows.

Two cases for the direction of evolution occurs. First, one can assume an allocative tradeoff between *r*_1_(*z*) and *α*_12_, i.e., the sign of d*r*_1_(*z*)/d*z /* d*α*_12_(*z*)/d*z <* 0 is negative. In such a case, the direction of evolution is determined by the strength of this trade-off and by the geometric of coexistence given by the angles Ω and *η*_2_, as well as the length of vectors **r** and **v**_2_. In the second case, where no trade-off is assumed, the signs of d*r*_1_(*z*)/d*z* and d*α*_12_(*z*)/d*z* are the same, and therefore, the direction of evolution is determined by their sign.

The effects of evolution on the different metrics of coexistence are given by:

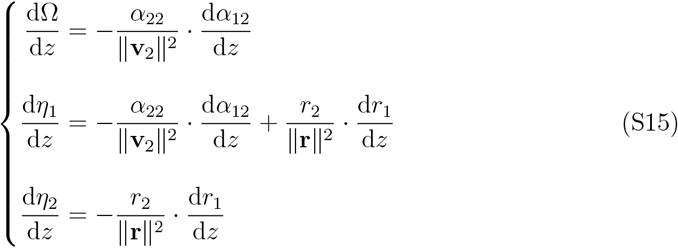

Note that dΩ/d*z* = d*η*_1_/d*z* + d*η*_2_/d*z*, which is coherent as, Ω = *η*_1_ + *η*_2_.

To easily relate these three equations to figure S4, we can rewrite them to make apparent the changes in *ω*(*z*_*m*_, *z*), Ω, *η*_1_, and *η*_2_ relative to the change in *α*_12_. This can also be considered equivalent to considering *z* = *α*_12_ and, consequently, *r*_1_ being a function of *α*_12_. This leads to

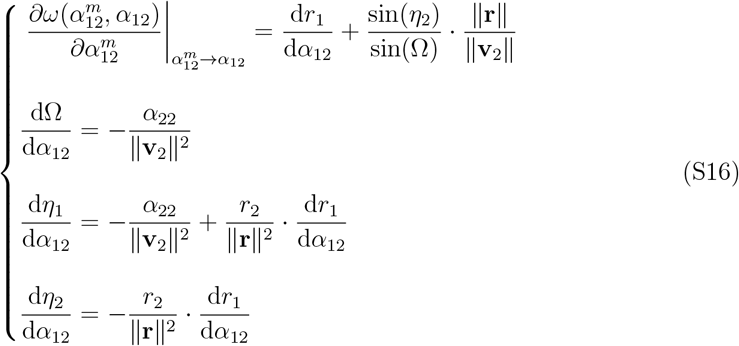

The first equation shows that *α*_12_ is selected if and only if sin(*η*_2_)*/* sin(Ω)·∥**r**∥*/*∥**v**_2_∥ *>* − d*r*_1_/d*α*_12_. That is, the geometry of coexistence (the second term of this equation) determines a threshold for selection and counter-selection. The second equation demonstrates that the evolution of *α*_12_ and Ω are in the same direction as we can assume *α*_22_ *<* 0 (intraspecific competition), which is coherent with figure S4. The fourth equation demonstrates that whether the evolution of *α*_11_ and *η*_1_ are in the same or opposite direction is determined by the sign of *r*_2_ · d*r*_1_/d*α*_11_. As we assume d*r*_1_/d*α*_11_ *<* 0, then if *r*_2_ *>* 0 evolution is in the opposite direction, as illustrated in figure S4, while it is in the same direction if *r*_2_ *<* 0. The reader can be geometrically convinced of the latter case by imagining the vector **r** to be located in the fourth orthant on panels B and D. Finally, the last equation shows that the determination of the direction of evolution of *η*_1_ is more tricky, as already shown in figure S4, as it will depend on the signs of *r*_2_ *>* 0, *α*_21_ *>* 0, and d*r*_1_(*z*)/d*z /* d*α*_11_(*z*)/d*z*, as well as on the amplitude of the vectors **r** and **v**_1_.

## S4 Evolution of a phenotype affecting *α*_11_ and *α*_12_

Here, we assume that the trait *z* of species 1, which is under selection, impacts both the intraspecific interaction *α*_11_ (intraspecific competition) and the interspecific interaction *α*_12_(*z*) (*per capita* effect of species 2 on the per capita growth rate of species 1). Such a situation can, for instance, relate to when prey invest in defence, i.e., a reduction of the negative impact of the predator that translates to an increase in *α*_12_(*z*), at the cost of an increase in its intraspecific competition (Agrawal et al., 2012) (i.e. ecological costs). Figure S5 shows the graphical analysis of the consequences of evolution on the geometry of coexistence for the four possible combinations of interactions. In this figure, we assume that evolution decreases the negative effect of species 2 on species 1 (panels A and D), or decreases the positive effect (panels B and C), but at the cost of an increase in the intraspecific competition, i.e., evolution results in a simultaneous decrease in *α*_11_ (that therefore becomes more negative; blue arrow on vector **v**_2_) and increase in *α*_12_ (that therefore becomes less negative or more positive; blue arrow on vector **v**_1_). Other cases can be obtained simply be first choosing the appropriate direction of the red and blue arrows, and then deducing the direction of the black arrows. The figure shows that the effects of evolution on both resistance angles *η*_1_ and *η*_2_ is uniquely determined. The evolution of *η*_1_ is always in the same direction as of *α*_12_ (all panels), while the direction of *η*_2_ evolution is opposite if species 1 as a positive effect on species 2 (panels B and D). The direction of Ω is also uniquely determined and goes in the same direction as *α*_12_, when, species 1 negatively impacts species 2 (panels A and C). It is more difficult to determine graphically when the effect of species 1 on species 2 is positive, as both vectors **v**_1_ and **v**_2_ turn in the same direction. In such instances, as mathematically demonstrated below, the impact of evolution on Ω is determined by the combination of the trade-off strength, d*α*_12_(*z*)/d*z /* d*α*_11_(*z*)/d*z*, and the vectors **v**_1_ and **v**_2_.

Following the same rationale as in section S2, the invasion fitness of a rare mutant of trait *z*_*m*_ in a resident population of trait *z* is given by

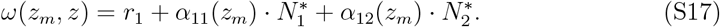

**Figure S5:**
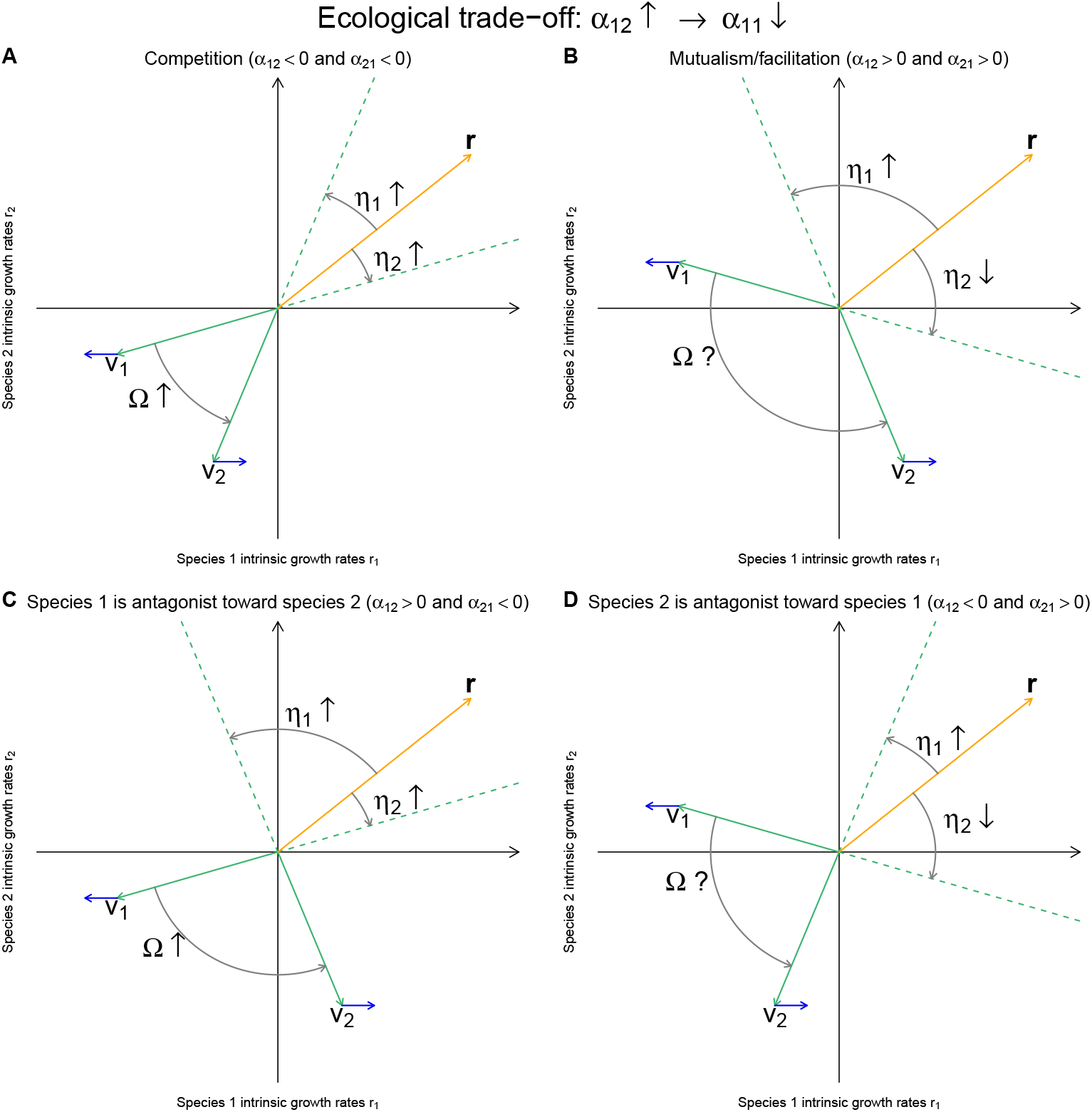
Effect of evolution of species 1 on the structural metrics in the case of an allocative trade-off between *α*_11_ and *α*_12_. Here, we assume that evolution relaxes the negative effect of species 2 on species 1 (evolution toward better defences), or increase the positive effect, at the cost of an increase in the intraspecific competition. Other cases can be obtained by simply setting accordingly the red and blue arrows, and then, deducing the direction of the black arrows.

The direction of trait evolution is then given by the selection gradient

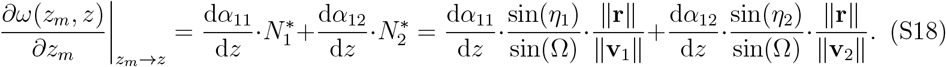

Two cases for the direction of evolution occur. First, one can assume an allocative tradeoff between *α*_11_ and *α*_12_, i.e., the sign of d*α*_12_(*z*)/d*z /* d*α*_11_(*z*)/d*z <* 0 is negative. In such case, the direction of evolution is determined the strength of this trade-off and the geometric of coexistence given by the angles Ω, *η*_1_, and *η*_2_, as well as the length of the vectors **r, v**_1_, and **v**_2_. In the second case, where no trade-off is assumed, the signs of d*α*_11_(*z*)/d*z* and d*α*_12_(*z*)/d*z* are the same, and therefore, the direction of evolution is determined by their sign.

The effects of evolution on the different metrics of coexistence are given by

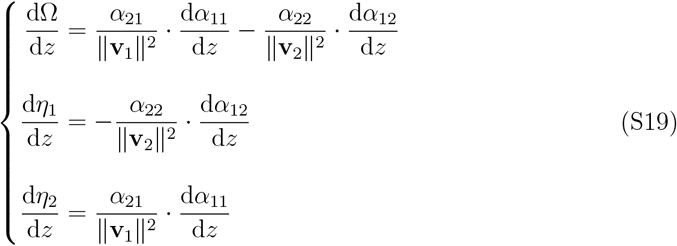

Note that dΩ/d*z* = d*η*_1_/d*z* + d*η*_2_/d*z*, which is coherent as, Ω = *η*_1_ + *η*_2_.

To easily relate these three equations to figure S5, we can rewrite them to make apparent the changes in *ω*(*z*_*m*_, *z*), Ω, *η*_1_, and *η*_2_ relative to the change in *α*_12_. This can also be considered equivalent to considering *z* = *α*_12_ and, consequently, *α*_11_ being a function of *α*_12_. This leads to

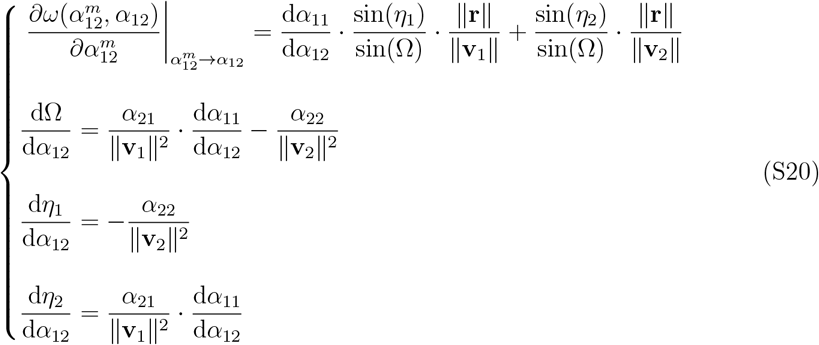

The first equation shows that *α*_12_ is selected if and only if 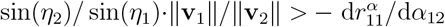. That is, the geometry of coexistence (the second term of this equation) determines a threshold for selection and counter-selection. The third equation demonstrates that the evolution of *α*_12_ and *η*_1_ are in the same direction as we can assume *α*_22_ *<* 0 (intraspecific competition), which is coherent with figure S5. The last equation demonstrates that whether the evolution of *α*_12_ and *η*_1_ are in the same or opposite direction is determined by the sign of *α*_21_ · d*α*_12_/d*α*_11_. As we assume d*α*_12_/d*α*_11_ *<* 0, then if *α*_21_ *>* 0 evolution is in the opposite direction, as illustrated in figure S5, while it is in the same direction if *α*_21_ *<* 0. Finally, the second equation shows that only in the cases of *α*_21_ · d*α*_12_(*z*)/d*z /* d*α*_11_(*z*)/d*z <* 0, the direction of Ω evolution is uniquely determined and in the same direction as *α*_12_. Otherwise, the direction of evolution is also dictated by the strength of the trade-off, d*α*_12_(*z*)/d*z /* d*α*_11_(*z*)/d*z* and the sign of *α*_21_.

## S5 Evolution of a phenotype affecting *r*_1_, *α*_12_, and *α*_21_

Empirical examples in figures 2 and 3 assume that the trait *z* of species 1 under selection impacts the two interspecific interactions *α*_12_ and *α*_21_, as well as the intrinsic growth rate *r*_1_. In the following, we derive the analytic computations for the invasion gradient and the evolution of metrics of coexistence, following the same rationale as in sections S2, S3, and S4.

The invasion fitness of a rare mutant of trait *z*_*m*_ in a resident population of trait *z* is given by

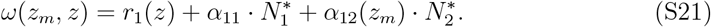

We remark that, as the interaction *α*_21_ does not intervene in the per capita growth rate of the rare mutant, this is the exact same expression as in section S3. Then, the direction of trait evolution is then given by the selection gradient

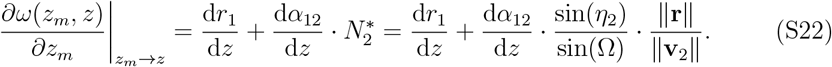

The effects of evolution on the different metrics of coexistence are given by

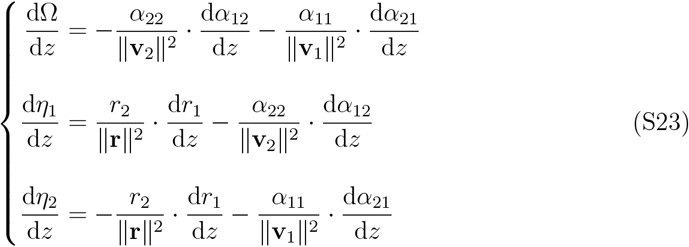

Note that dΩ/d*z* = d*η*_1_/d*z* + d*η*_2_/d*z*, which is coherent as, Ω = *η*_1_ + *η*_2_.

To relate these three equations to figures 2 and 3, we can rewrite them to make apparent the changes in *ω*(*z*_*m*_, *z*), Ω, *η*_1_, and *η*_2_ relative to the change in *α*_12_. This can also be considered equivalent to considering *z* = *α*_12_ and, consequently, *α*_21_ and *r*_1_ being a function of *α*_12_. This leads to

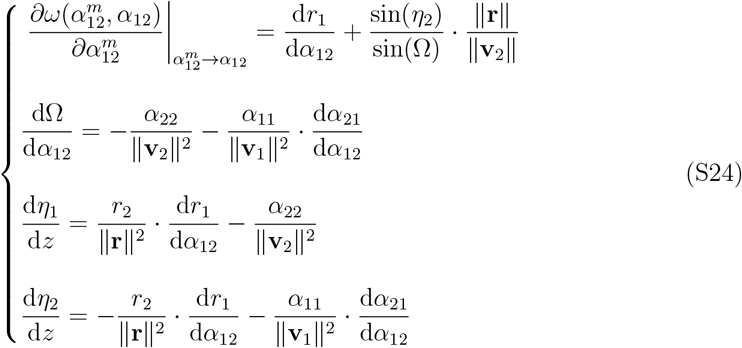

